# Comments on the recent crystal structure of TsaBDE complex of bacterial t^6^A biosynthesis system and its significance for understanding TC-AMP processing

**DOI:** 10.1101/563171

**Authors:** Boguslaw Stec

**Author notes:** 3902 Broadlawn St. San Diego, CA 92111.

## Abstract

The N(6)-threonylcarbamoyl adenosine (t^6^A) modification at position 37 of a tRNA of the anticodon loop is universal and central to the translational fidelity of all known organisms. The ternary complex of TsaBDE is the central and essential workstation for t^6^A biosynthesis in bacteria. The recently published crystal structure of *Thermotoga maritima* (*T.maritima*) TsaBDE complex (Missoury *et al.*, 2018) has ~15% incorrectly-placed, misplaced/mistraced, or missing residues. These structural errors have precipitated incorrect conclusions about the disordering of the active site and inferred action of the TsaE element. In this report, we rectify the published structural model of the *T.maritima* TsaBDE complex. In stark contrast, a corrected structural model of TsaBDE shows that both active sites of the TsaD element are fully occupied with threonylcarbamoyladenosine (TC-AMP), an unstable intermediate chemical moiety of the t^6^A biosynthesis pathway. This observation has profound implications for understanding the funneling of intermediates in the t^6^A pathway and also in helping to elucidate tRNA binding modes. Based on the structural details described in here we propose a unifying principle for binding the tRNA to the TsaD subunit of the complex which is universally required in all known t^6^A modification pathways.

## Introduction

The biosynthesis of N(6)-threonylcarbamoyladenosine (t^6^A) is a complex multistep process in bacteria that involves a family of proteins: TsaB, TsaC/TsaC2, TsaD, and TsaE (Deutsch *et al.*, 2012, Luthra *et al.*, 2018). It is well-demonstrated that the TsaC/C2 protein catalyzes the condensation of Thr-CO_2_/CO_3_-ATP to obtain an unstable intermediate threonylcarbamoyl-AMP (TC-AMP) (Lauhon, 2012). It is believed, but not directly experimentally demonstrated, that the direct interaction of TsaC/C2 with the TsaD component is responsible for a transfer of this intermediate into the active site of the TsaD where, in conjunction with TsaB, a catalytic transfer of threonylcarbamyl (TC) moiety is carried onto N6 of adenosine 37 of the incoming tRNA molecule (Luthra *et al.*, 2018, Missoury *et al.*, 2018). This belief is deeply rooted in a conviction that there is an insufficient concentration of the freely available unstable intermediate that would persist under conditions of intense stress which usually stimulates rapid tRNA synthesis. Additionally, the belief is supported by the existence of a highly structurally homologous enzyme, TobZ, which has dual functionality corresponding to a fused pair of TsaC/TsaD (Parthier *et al.*, 2012).

What was also not directly experimentally demonstrated, is the direct competition between the binding of tRNA and the TsaC/C2 component which would be expected if the mode of binding was overlapping. The latest attempt at showing a direct interaction between TsaC/C2 and TsaD with any of the stable complexes of TsaBD was inconclusive (Luthra *et al.*, 2018). What was demonstrated is the direct competition of tRNA with the TsaE component, but surprisingly in a different concentration range from that at which the TsaE stimulates the tRNA binding (Luthra *et al.*, 2018). More experimental evidence is needed to prove that a direct interaction between the TsaC/C2 and TsaD subunits is required for the successful production of the (t^6^A) modification.

Eukaryotic systems employ catalytic core proteins TsaC/TsaC2 and Kae1, an analog to TsaD (El Yacoubi *et al.*, 2009, Srinivasan *et al.*, 2011), but use a variety of additional proteins as regulators of the complex (Gon7/Pcc/Bud32/Cgi21) (Srinivasan *et al.*, 2011, Perrochia, Guetta*, et al.*, 2013, Zhang, Collinet, Graille*, et al.*, 2015). High structural and sequence homology of the central proteins such as TsaD/Kae1/Qri7 (Thiaville *et al.*, 2014) suggests a similar mode of operation as well as a similar mode of tRNA binding. Therefore, finding out a precise position for the bound substrate would offer a significant advantage in an attempt to model the active tertiary complex TsaD-TC-AMP-tRNA. An initial tentative model of the docked TC-AMP was published by the Missoury *et al* (Missoury *et al.*, 2018). To further advance the knowledge of the details of the catalytic transfer of the TC to N6A, the Tilbeurgh team obtained a complex of TsaBDE from *T. maritima* (Missoury *et al.*, 2018, Luthra *et al.*, 2018).

The structure of the complex was solved and refined at a 3.1Å resolution (Missoury *et al.*, 2018). A closer inspection of the model showed many internal inconsistencies and structural errors (Fig 1). The main inconsistency was a noticeable asymmetry in the otherwise symmetric model, which combined with significant departures from the previously obtained crystallographic models of individual components, raises doubts about the reliability of this complex and the conclusions drawn from the observations. Also, the mechanistic implications suggested by the authors concerning the role of the TsaE element cannot possibly be correct when most of the erroneously-refined fragments are in its vicinity.

**Figure 1:**
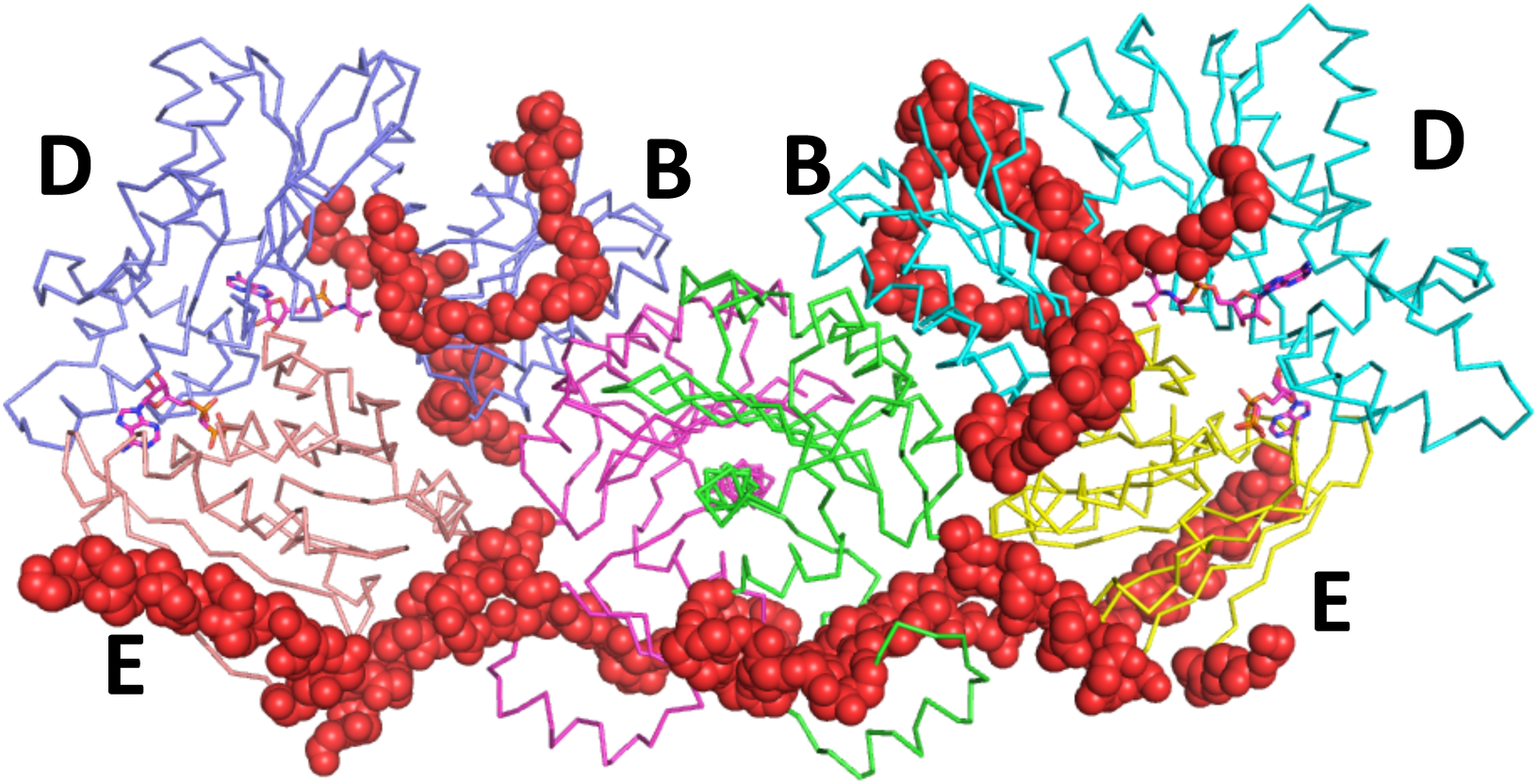
The general view of the final refined B_2_D_2_E_2_ complex in C-alpha representation. Different subunits are represented in different colors. The substrates are represented by purple stick models. The fragments of the model that were wrongly refined (one-, two-residue shifts, misplaced or missing) are represented in red spheres. These encompass approximately 15% of the structure and closely surround the active sites of TsaD.

In order to resolve these issues, the model and the data (Missoury *et al.*, 2018) were retrieved from PDB and the model was re-refined against the original data. The corrections to the TsaBDE complex described here have significant implications for understanding the entire pathway. The corrected model shows relatively well organized active sites of the TsaD and TsaE and reveals both TsaD active sites fully occupied with an unstable substrate TC-AMP. This result contradicts the notions of the authors that the main role of the TsaE element is to disorder the active site. The presence of the TC-AMP sheds light onto the functioning of the entire pathway. It allows one to postulate the chemical mechanism for TC transfer to the tRNA and imposes the limitation on the tRNA binding mode. We propose a tentative model of the tRNA binding to the TsaD that offers a transferability of consensus interaction across different kingdoms, species, and organelles as well as offers a universal unifying mechanism of action of the TsaD element and its analogs (Thiaville *et al.*, 2014).

The corrected structure in combination with previous biochemical results sheds additional light on the role of the TsaE component in regulation of the TsaBD dimer and conjunction of the TsaD component with tRNA.

## Materials and methods

### Crystallography

The protein coordinates and diffraction data extracted from PDB were imported to CCP4. The mtz file was created using F2mtz (import) option with 5% of data in the test set. This mtz file together with the PDB file was used for the refinement in Refmac (Vagin *et al.*, 2004). The comparison of the statistics between the initial and final model is in Supplementary Table S1. A similar protocol was applied to the model of Qri7 (3WUH) (Tominaga *et al.*, 2014) to clarify the structural details of the missing fragments. We also retrieved a family of structures describing the details of the Kae1 complex (Mao *et al.*, 2008) which, without further refinement, were used in modeling. (PDB structures 3ENH, 3EN9, 4WW5, 4WXA) (Srinivasan *et al.*, 2011, Perrochia, Guetta*, et al.*, 2013, Zhang, Collinet, Graille*, et al.*, 2015, Mao *et al.*, 2008).

### Modeling

The refined models of the Qri7 dimer (Tominaga *et al.*, 2014), B_2_D_2_E_2_ complex (Missoury *et al.*, 2018), and a complete assembled model of the Kae1 complex containing Gon7, Pcc1, Kae1, Bud32, and Cgi121 (Mao *et al.*, 2008) were used in constrained docking experiments to address possible tRNA binding modes. The tRNA model was taken from the crystal structure of tRNA in complex with a charging synthase (1qf6) (Sankaranarayanan *et al.*, 1999). First, we inspected the models for completeness and possible conformational mobility by re-refining individual PDB structures and inspecting the distributions of the temperature factors in resulting models.

All structures used in modeling were thoroughly or lightly refined using Refmac (Vagin *et al.*, 2004) included in the CCP4 release 7.0. The TsaBDE complex (Missoury *et al.*, 2018) was thoroughly refined resulting in significant improvements described below. Qri7 (Tominaga *et al.*, 2014) was moderately refined to fill in missing pieces of the backbone, while elements of the Kae1 complex (Mao *et al.*, 2008) (3ENH, 3EN9), were only very lightly refined to improve stereochemistry.

### Docking

Docking was performed with multiple docking programs (Tuszynska *et al.*, 2015, van Zundert *et al.*, 2016, Morris *et al.*, 2009) (Haddock, Autodock, NP-Dock) that produced similar results, but without clear distinctions between the clusters. Haddock (van Zundert *et al.*, 2016) and Autodock (Morris *et al.*, 2009) lead to very similar results, but with more diverse cluster distribution than results of NP-Dock (Tuszynska *et al.*, 2015) which are discussed here. In order to perform docking experiments on the protein models that were newly refined or extracted from the RCSB-PDB database (as described above), we determined tentative catalytic sites and established a reaction sphere surrounding the amide bond in TC-AMP. In all docking experiments, we used the tRNA model (1QF6 (Sankaranarayanan *et al.*, 1999)) with a distance constraint that placed N6 of A_37_ within the active site sphere of a protein. The resulting 30 clusters were ranked by pseudo energy and visually inspected. The clusters with the highest pseudo energy and not overlapping with the crystallographically determined positions of the protein binding partners (TsaC/C2 or TsaE) were selected for analysis.

## Results and Discussion

### Refinement of the TsaBDE complex from T. maritima

The model of *T. maritima* Tsa proteins BDE 2:2:2 complex was retrieved from PDB (6FPE) (Missoury *et al.*, 2018) together with the diffraction data associated with it. After several initial cycles of refinement using Refmac 5.8, the initial electron density maps indicated many problems with the presence of excessive positive and some negative electron density. The existing models for all components (TsaB/D/E) were compared to the previously solved and deposited models, such as 3ZEU (Nichols *et al.*, 2013) for the TsaBD complex, 2A6A (Xu *et al.*, 2010) for the TsaB dimer, or 1HTW for TsaE (Teplyakov *et al.*, 2002). Careful examination of the individual proteins showed numerous tracing errors as well as untraced elements of the model. The final model contains more than 50 newly refined residues and more than 190 of existing residues were relocated to new positions. This accounts for more than 15% of the structure that was displaced or not refined. The newly refined elements improved the standard R factor (0.196 from original 0.223) while leaving the Rfree at almost the same level (0.298 versus 0.294 for the original model). The stereochemistry of the model has been improved by correcting more than 30% of residues in reference to their preferred conformers. General improvements allowed for obtaining a much better fit to the electron density and discovery of fully occupied threonylcarabamoyl-AMP (TC-AMP) in both active sites of the TsaD subunits (Fig. 2).

**Figure 2:**
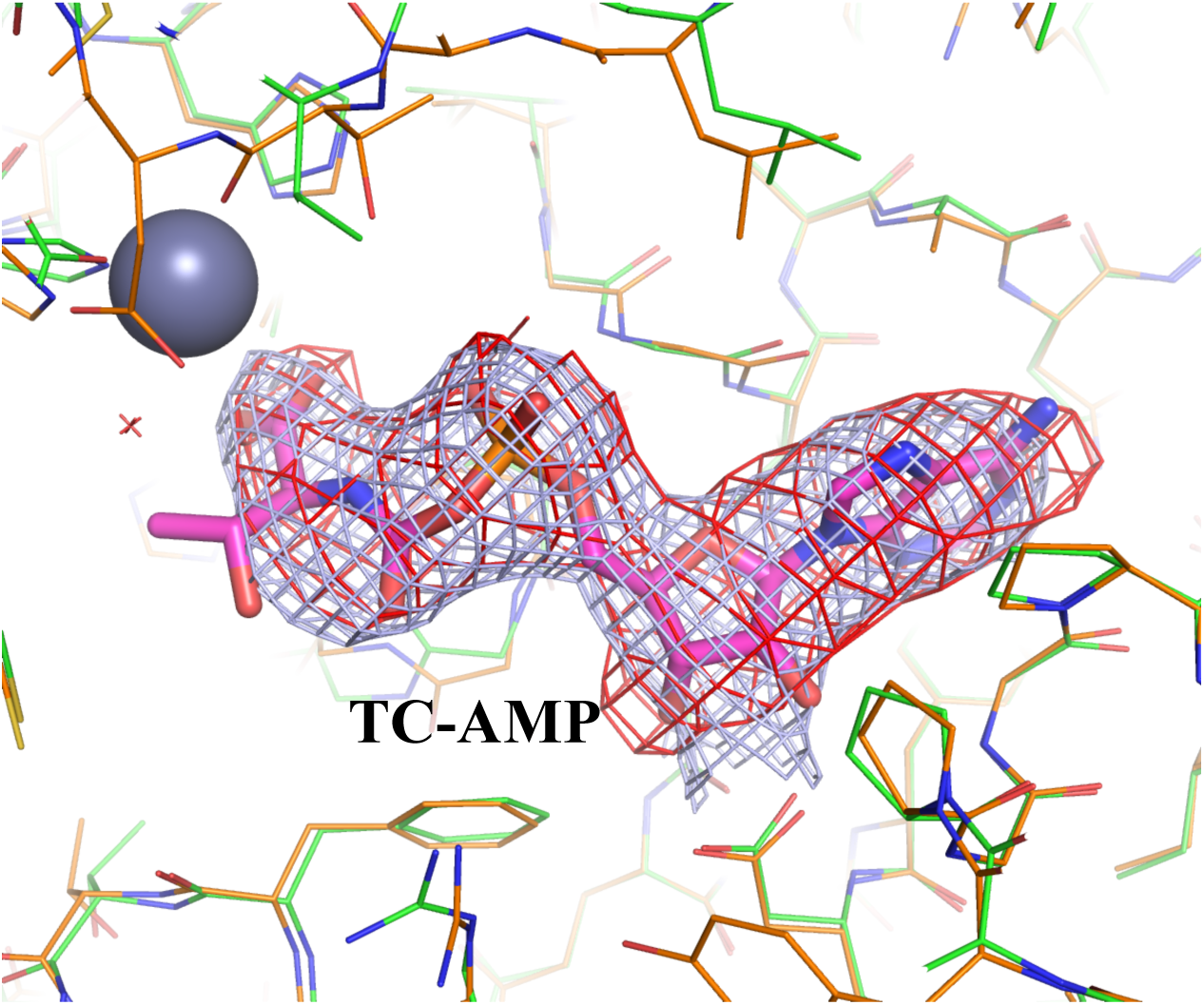
A Fully occupied model of the TC-AMP molecule in the active site of the TsaD component. The omit map in red (3σ) and the 2Fo-Fc ED map passed with the final refined model contoured at a 1.3σ level in bluish-gray, are presented. Final refined model is in orange while the initial model is in green. The large gray sphere represents the zinc atom refined with 40% occupancy.

In both TsaB type subunits, the two regions 112-137 and 190-206 were the most notable. The first region (112-137) shows a single amino-acid shift as compared with the final model and the previously published structure. The second region (190-206) shows misplaced and unrefined residues where the difference maps allowed for more complete modeling. These errors were clearly identifiable when the initial structure was compared to the published structure of the B dimer (2A6A) (Xu *et al.*, 2010).

TsaD showed larger errors with a surprising asymmetry between both subunits, particularly in the region, 294-327. A comparison of both subunits shows a shift by one residue in the region 297-309. The comparison of both models superposed and covered with the final 2Fo-Fc and with difference maps is shown in supplemental Fig S1. Additionally, when compared to the final model both subunits showed a shift of two residues in the final 309-327. Therefore, an initial model had two residues 325-327 missing from the model. Both TsaD subunits were missing residues 31-49 which are one of the key conformational differences that can be attributed to binding of the TsaE element, as can be deduced by comparing it to homological fragments in other TsaD type models, like in 3ZEU (Nichols *et al.*, 2013).

The subunit TsaE refined relatively well but the quality of the electron density was not as high as for other subunits. However, at closer inspection some additional unmodeled electron density was located near the N-terminus and it appeared that the entire C-terminal helix was not optimally rotated. The closer analysis of the hydrophobic moments combined with the comparison to the homologous domains suggested a single position rotation. The correct placement of the helix allowed for more residues to be modeled at the C-terminus, resulting in new positions for residues 145-162.

Overall improvements to the model, particularly around the active site, resulted in the appearance of the electron difference densities of a large ATP-like molecule. In the original model (Missoury *et al.*, 2018), only one site was partially modeled with a glycerol and short PEG. The final model was refined with full occupancy TC-AMP at both active sites of subunit TsaD. After modelling the substrate TC-AMP, some residual electron density suggested the presence of partially occupied metal ions. Zinc ions were refined with 40% occupancy at both active sites. The only fragments at both active sites that were weakly defined by electron density were residues 291-294. These residues show more extended conformation than the same region in homologous proteins that attains alpha helical conformation. It appears this transition accommodates a more twisted/open conformation of the substrate binding domain of the TsaD as visualized in Supplemental FigS2. This domain shows a 17° rotation as compared to the reference structure of the *E. coli* TsaB-TsaD heterodimer (4ydu) (Zhang, Collinet, Perrochia*, et al.*, 2015).

This is the first observation by X-ray crystallography of the TC-AMP, which is unstable in aqueous solutions. The full occupancy of this moiety is quite an important observation as it has only 2.5 minutes of half-life time in solution (Lauhon, 2012). The presence of it in the crystal structure would suggest that this compound was protected by binding to the protein to survive the entire crystallization and structure determination process. The real burning question is about the origin of this compound. The original paper clearly presented the method of obtaining the samples of the proteins. Missoury *et al* (Missoury *et al.*, 2018) described the process as the production of active DBBD dimers *in vivo* and purifying the complex. This clearly suggests that TC-AMP in vivo synthesized by native *E.coli* TsaC could have been captured by DBBD dimer, survived the purification and crystallization process, and was detected in the final re-refined model. Another interesting fact is the decorrelation of the occupancy of this substrate from the presence of the metal ion. Fractional occupancy of the metal ion suggests that its presence is indispensable for the transfer of the TC moiety to the A37, but not necessary for binding of the TC-AMP molecule. That creates a very interesting situation in which binding of the metal ion that is coordinated with the presence of ATP and its hydrolysis by the TsaE element controls the activity of the entire complex.

### Main structural changes to the DBBD dimer caused by binding of the TsaE element

Two biochemical findings that are important for understanding the role of TsaE in controlling the activity of BD or DBBD complexes is that it competes with tRNA, but also that, at the same time, it primes the binding of tRNA to the TsaBD complex (Luthra *et al.*, 2018). It is important how these two apparently opposing roles can be accomplished by the same molecule. Luthra *et al*. (Luthra *et al.*, 2018) demonstrated that this general control is exerted by the ATPase activity of the E component. The symmetric dimer of BDE presented by Missoury *et al.* (Missoury *et al.*, 2018) and amended by this report clearly shows some interesting features that can help to understand this regulation. In Qri7 (Wan *et al.*, 2013), the TsaD element forms a TsaD homodimer so only an internal mobility of a dimer can control tRNA binding. Since it is a symmetric dimer it cannot process two tRNA at the same time, due to interference with each other. The TsaBD heterodimer that is formed in bacteria creates a new possibility for external control. The BD interface is similar to a D-D homodimer interface in Qri7 in design and follows a logic of a four-helical bundle. Therefore, the refined structure gives an insight into the intimate relations of TsaE and TsaD as well as their mutual interactions that results in conformational changes to both.

The largest conformational divergence shown when the TsaBD heterodimer from *E. coli* (4YDU) (Zhang, Collinet, Perrochia, *et al.*, 2015) is superposed with the *T. maritima* TsaBD complex is in the region of amino acids of 30-40 of TsaD. This conformational change is depicted in the Fig. S2. This almost 90 degree rotation is coupled also with a significant opening (around 17°) of the domains of the D subunit. The *E. coli* structure shows a much narrower cleft at the apex of which the substrate is binding. Presumably limiting the space in this cleft is also needed for binding of tRNA and transfer of the TC group to A37. This is an important observation because it shows how both subunits are primed for certain action by a mutual relationship. In the figure, the TsaE subunit clearly penetrates and clashes with the 30-40 fragment of the TsaD subunit in the *E. coli* model which results in a 90° rotation of this fragment in *T. maritima* model. Additionally, binding of TsaE opens up the binding cleft of the TsaD subunit by displacing the nucleotide binding domain of TsaD, thus facilitating leaving of any remaining groups at the active site in the post-catalytic complex.

### Modeling of the tRNA binding to the TasD-like subunits of the t^6^A_37_ eukaryotic and prokaryotic complexes

The detection of TC-AMP at the active site of the TsaD component raises some very interesting possibilities. The mechanism of the transfer of threonylcarbamate moiety can be postulated as a nucleophilic attack of the activated N6 nitrogen on the bridging carbon in TC-AMP and formation of the tetrahedral intermediate. The unstable tetrahedral intermediate later collapses to the product, driven by an AMP leaving group. Therefore, the location of the flat face of the pseudo-peptide amide bond in TC-AMP defines a location of the approaching N6 of the adenosine ring with high accuracy (Fig. 3A). The most likely location for a deprotonated N6 approaching face to face with TC-AMP must be a sphere with an approximate diameter of 3Å for an effective transfer. Applying the constraint described above (N6 within the sphere), the comprehensive docking experiment was performed on three representative complexes producing *t^6^A*_37_ in mitochondria, in yeast, and *T. maritima*. The *E. coli* and *Bacillus subtilis* complexes can be understood as half complex of *T. maritima* and therefore they were not used despite being available in PDB.

**Figure 3.**
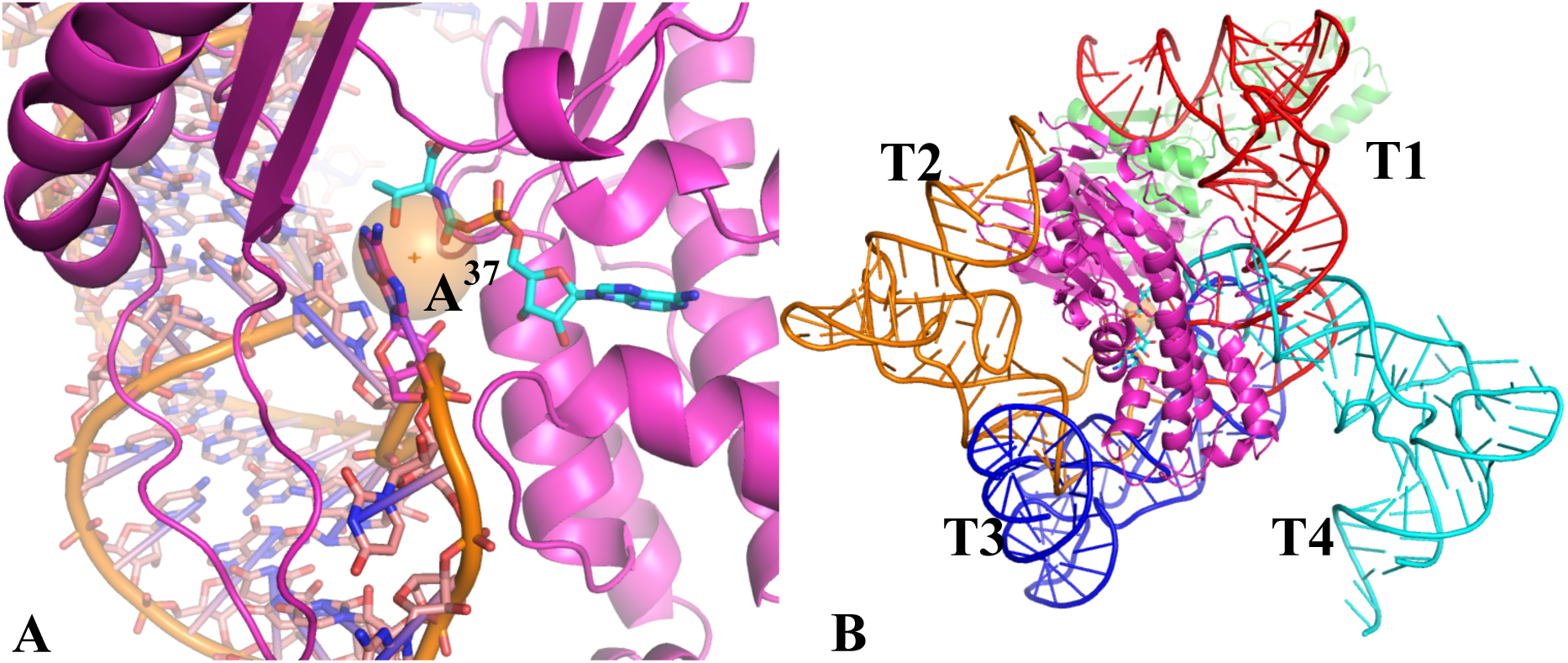
The docking of the tRNA to the TsaD subunit (and homologs). A) The most likely pose with a sphere drawn next to the TC-AMP representing the constraint between a protein and the tRNA. B) Four representative clusters that fulfill the constraint. The most likely pose (T1) is in red. Remaining poses are eliminated by the requirement for auxiliary protein binding (T2, T3) or insufficient binding energy (weak stabilization of the complex – T4).

The dockings were carried out with the refined version of the DBBD dimer of the *T.maritima* complex described in this report (with TsaE omitted), the modestly refined Qri7 dimer, and the reconstituted model of the yeast Kae1 complex described before. All dockings were performed with the tRNA model extracted from PDB (1QF6)(Sankaranarayanan *et al.*, 1999), a model of the tRNA complex with a threonyl tRNA synthetase, using the constraint described above. The web service NPDock (Tuszynska *et al.*, 2015) was used. As a standard result, the program produces 30 clusters of individual solutions ranked by pseudo-energy. In general, the results obtained can be classified as mostly four dominant poses. These poses (shown in Fig 3B) allow for A_37_ to be positioned properly to allow for the transfer of the TC moiety and provide sufficient binding energy with the protein complex. The rules that we adopted for selecting the best solution were: (1) the highest position on the rank pseudo-energy list, (2) feasible geometrical positions for the transfer of the TC moiety, and (3) non-overlapping positions with the postulated binding sites for auxiliary proteins (especially TsaC/C2 or TsaE) needed for production and translocation of TC-AMP. Out of the four best clusters (T 1-4), only one satisfied all the conditions (T1).

The best pseudo-energy results were obtained for Qri7, and while the TM model produced intermediate results, the least distinct solution was obtained for the Kae1 complex. However, the best solution for all these complexes was in full agreement with each other and established a consensus binding pose that was utilized by the entire family, regardless of the presence of the auxiliary proteins (Fig 4). This result suggests the possible pathway of increasing regulatory control by the gradual addition of individual proteins to the core built around the TsaD catalytic subunit. This conclusion is supported with recent observations that TsaE component in the TsaBDE complex (Luthra *et al.*, 2018, Zhang, Collinet, Perrochia, *et al.*, 2015), as well as Bud32 in the KEOPS complex (Perrochia, Crozat, *et al.*, 2013, Perrochia, Guetta, *et al.*, 2013), bind and process ATP that is located a the interface between D and neighboring subunit.

**Figure 4:**
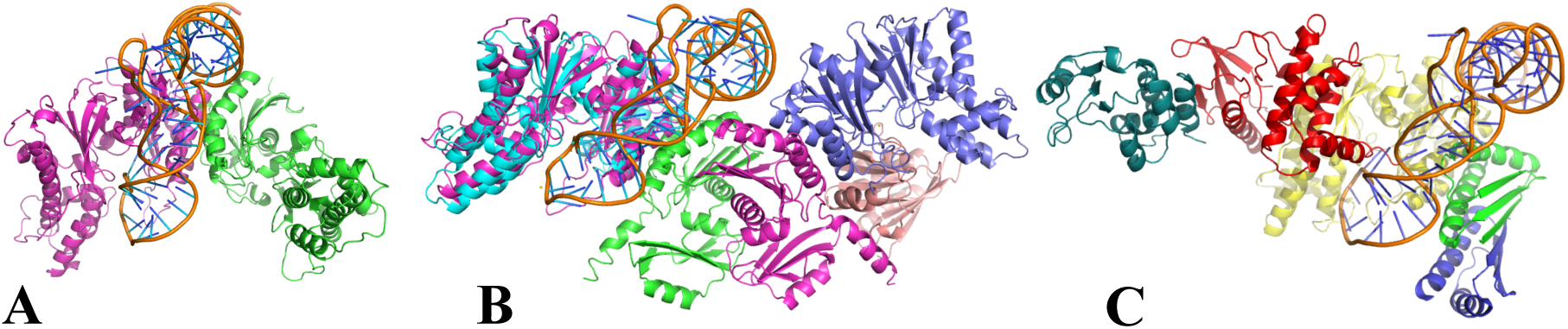
The predicted consensus poses for tRNA docking to the eukaryotic and prokaryotic complexes of t6A synthesis. (A) Qri7 homodimer with the docked tRNA. (B) *T. maritima* B_2_D_2_E_1_ + tRNA (TsaD in blue is superposed on the Qri7 monomer from (A) in purple). (C) Kae1 complex from the yeast with tRNA. The Kea1 subunit homologous to TsaD is in yellow.

The universality of the binding pose for tRNA proposed in this report raises a very interesting question. Why so vastly different complexes accomplish the same catalytic role of transferring the TC group to the A_37_. It appears that an attractive hypothesis is provided by an extension of an already described mechanism of increasing control of the TsaBD complex reactivity by binding of auxiliary proteins. Eukaryotic KEPOS complex is known to possess DNA binding capabilities as well as telomere editing activity (Liu *et al.*, 2018). This activity may require an increased ability to bind longer stretches of DNA. Such a binding would be difficult to accomplish providing a single Qri7-like dimer(Wan *et al.*, 2013). The binding of auxiliary proteins not only provides a control over the activity of the complex but apparently also provides an increased capability for creating additional functions, like DNA binding. It was recently shown that the dimerization of the Kae1 complex through PCC1 is not necessary for the tRNA activity (Perrochia, Guetta, *et al.*, 2013), while this dimerization event may be required for DNA telomerase activity (Liu *et al.*, 2018).

## Conclusions

In this report we not only provided a corrected model for the *T. maritima* TsaBDE complex of the t^6^A_37_ tRNA modification pathway, but also described two significant consequences of such a correction. We detected and described the unstable substrate TC-AMP whose position at the active site allowed for proposing a unified mode of tRNA binding to such complexes. Both findings are in full agreement with previously described biochemical results (Luthra *et al.*, 2018) and provide a constructive platform for future molecular understanding of this important biological process.

## Supplementary material / Appendix

**Supplementary Table S1.** Refinement statistics of the original and re-refined model of *T. maritima* TsaB-TsaD-TsaE complex in B2D2E2 composition and Qri7 from yeast

| Protein | *T. maritima* B<sub>2</sub>D<sub>2</sub>E<sub>2</sub> complex |  | Qri7 |  |
| --- | --- | --- | --- | --- |
| PDB code | Original model 6FPE | Re-refined model 6NAK | Original model 3WUH | Re-refined model 6NBJ |
| Space group | P212121 | P212121 | P4332 | P4332 |
| Unit cell dimensions |  |  |  |  |
| a (Å ) | 84.31 | 84.31 | 180.3 | 180.3 |
| b (Å ) | 113.99 | 113.99 | 180.3 | 180.3 |
| c (Å ) | 177.57 | 177.57 | 180.3 | 180.3 |
| Resolution (Å ) | 3.14 | 3.14 | 2.94 | 2.94 |
| Average redundancy | 4.56 | 4.56 | 3.8 | 3.8 |
| Average I/r (I) | 7.5 | 7.5 | 8.0 | 8.0 |
| Completeness (%) | 99.1 | 99.1 | 99.8 | 99.8 |
| Total R-merge (%) | 16.8 | 16.8 | 8.9 | 8.9 |
| No Refined Reflections | 30281 | 28747 | 21792 | 20688 |
| R-value (%) | 22.9 | 19.6 | 22.6 | 20.0 |
| Rfree-value (%) | 29.4 | 29.8 | 27.3 | 26.9 |
| No of residues | 1313 | 1470 | 676 | 752 |
| No of atoms | 10496 | 10993 | 5279 | 6084 |
| Rmsd, bond lengths (Å ) | 0.10 | 0.09 | 0.09 | 0.11 |
| Rmsd, bond angles (deg.) | 1.25 | 1.35 | 1.2 | 1.6 |
| Average B for subunitsB | 77, 68 | 87, 77 | NA | NA |
| Average B for subunitsD | 89, 76 | 101, 89 | 97, 99 | 103, 103 |
| Average B for subunitsE | 99, 73 | 111, 82 | NA | NA |
| Average B ligands at sitesD | 62 | 102, 93 | 113, 118 | 103, 108 |
| Average B ligands at sitesE | 96, 72 | 88, 75 | NA | NA |
| Ligands(D, E) | GOL, APC | TC-AMP,ATP | AMP | AMP, 4-C |
| Ramachandran (within) | 91.4 | 87.9 | 93.1 | 88.6 |
| Ramachandran (cond) | 7.5 | 8.8 | 5.5 | 8.2 |
| Ramachandran (outside) | 1.1 | 3.3 | 1.5 | 3.2 |

**Supplementary Figure S1:**
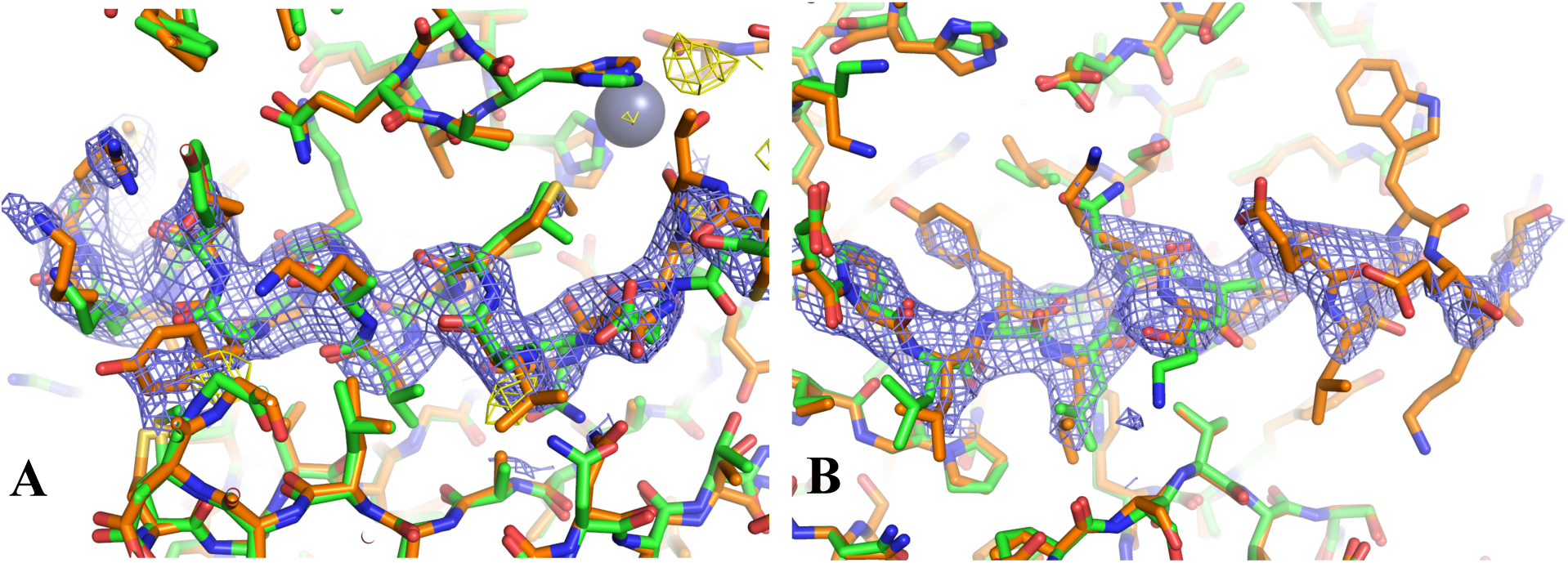
Electron density and model improvements during the refinement. The original model is represented in green stick models while the final refined model is in orange. The 2Fo-Fc electron density phased with the final model (in blue) is contoured at 1.2 σ level, while the difference Fo-Fc phased with the original model (in yellow) is contoured at 3σ. A) The fragment of the model around residues 280-300 in TsaD which clearly show a single residue shift, B) The additional residues and the improvement to the fragment of 200-230 in TsaB which shows mis-traced as well un-modelled residues.

**Supplementary Figure S2:**
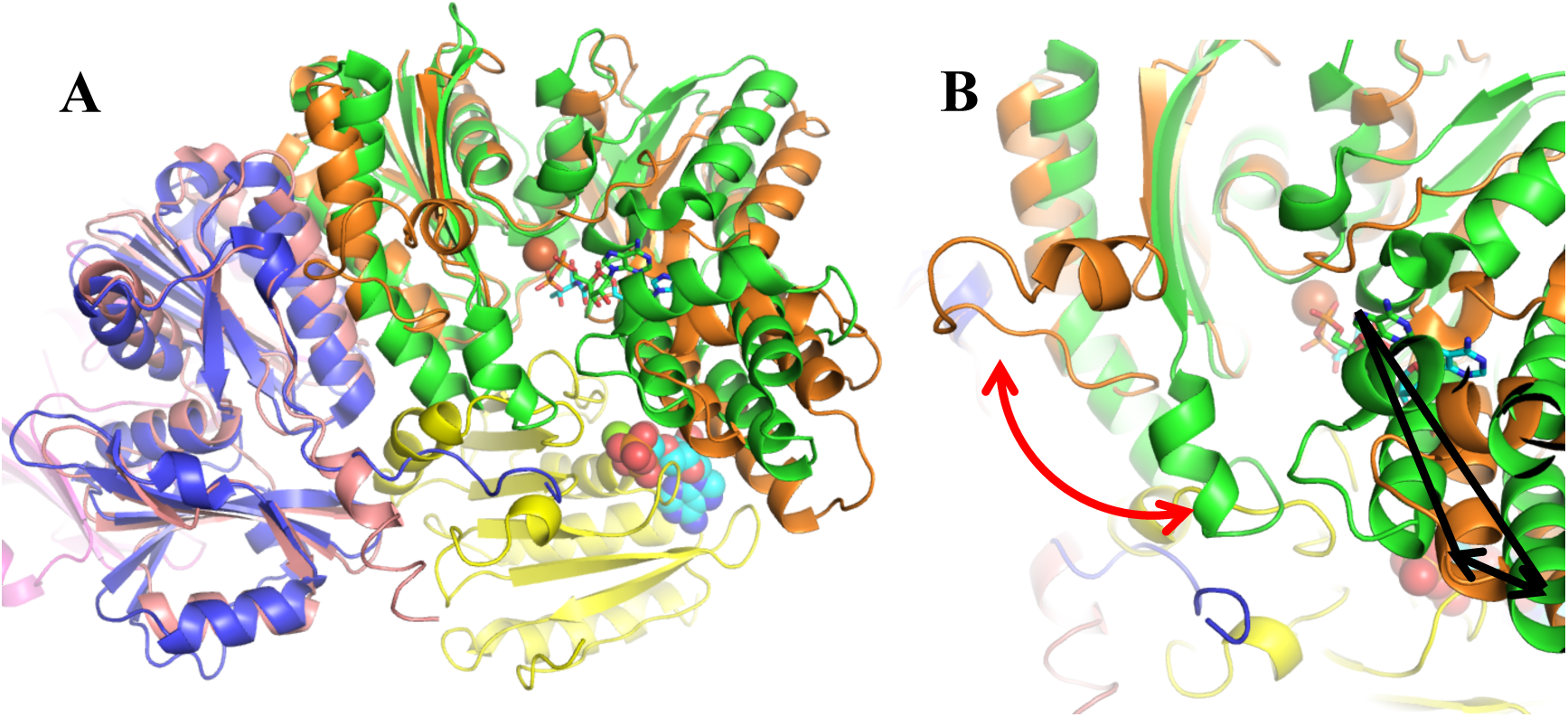
Superposition of the models of TsaBD dimer form *E. coli* (4ydu) and refined model of DBBD dimer from *T. maritima* with the TsaE bound at both sides. (A) The comparison of the BD dimer conformations (blue and green are *E. coli* BD subunits, pink and orange are *T. maritima* BD subunits while E subunit is in yellow), (B) A close-up of the conformational change of the 30-40 region in TsaD denoted by a red arrow. The domain movement is indicated by black arrow.

